# EpiCure (Epithelial Curation): a versatile and handy tool for curation of epithelial segmentation

**DOI:** 10.64898/2026.03.27.714683

**Authors:** Gaëlle Letort, Léo Valon, Arthur Michaut, Tom Cumming, Laura Xénard, Minh-Son Phan, Nicolas Dray, Curtis T Rueden, François Schweisguth, Jérôme Gros, Laure Bally-Cuif, Jean-Yves Tinevez, Romain Levayer

## Abstract

Investigating single-cell dynamics and morphology in tissues and embryos requires highly accurate quantitative analysis of microscopy images. Despite significant advances in the field of bioimage analysis, even the most sophisticated segmentation and tracking algorithms inevitably produce errors (e.g. : over segmentation, missing objects, miss-connected objects). Although error rate may be small, their propagation throughout a time-lapse sequence has catastrophic effects on the accuracy of tracking and extraction of single cell parameters. Extracting single cell temporal information in the context of tissue/embryo requires thus expert curation to identify and correct segmentation errors. In the movies commonly used in developmental biology and stem cell research, both the number of imaged cells and the duration of recording are large, making this manual correction task extremely time-consuming. This has now become a major bottleneck in the fields of development, stem cell biology and bioimage analysis. We present here EpiCure (Epithelial Curation), a versatile tool designed to streamline and accelerate manual curation of segmentation and tracking in 2D movies of large epithelial tissues. EpiCure uses temporal information and morphometric parameters to automatically identify segmentation and tracking errors and provides user-friendly tools to correct them. It focuses on ergonomics and offers several visualization options to help navigating in movies of tissue covering a large number of cells, speeding up the detection of errors and their curation. EpiCure is highly interoperable and supports input from a wide range of segmentation tools. It also includes multiple export filters, enabling seamless integration with downstream analysis pipelines. In this paper, using movies from several animal models, we highlight the importance of curating cell segmentation and tracking for accurate downstream analysis, and demonstrate how EpiCure helps the curation process for extracting accurate single cell dynamics and cellular events detection, making it faster and amenable on large dataset.

## Introduction

Extracting information about single cell behaviour, shape and signalling in the context of embryogenesis has become central to address core questions of developmental biology. The combination of novel sensors, fluorescent tracers and microscopy methods facilitate the generation of movies following the dynamics of tissues and organisms with single cell resolution over long time scales. However, their analysis via segmentation and tracking remains a strong bottleneck. This is still the case even in 2D epithelial tissues, which are among the most experimentally tractable systems in both embryonic and adult organisms, and where cells are tightly adherent and densely packed. Several powerful tools, often based on Deep Learning (DL), are now available for epithelia segmentation, *e.g.* CellPose (Stringer et al., 2021), EpySeg (Aigouy et al., 2020), Dissect for 3D (Merle et al., 2023) and DistNet2D (Ollion et al., 2024). These algorithms can readily perform tissue segmentation with only a small error rate, typically around 5-10% (Farrell et al., 2017; Funke et al., 2019; Mosaliganti et al., 2012; Turley et al., 2024; Wolny et al., 2020). This remains acceptable for a few applications, especially for single time point analysis, but becomes an issue for temporal dynamics. Indeed, the segmentation errors will propagate across time and strongly affect tracking accuracy (Bragantini et al., 2025; Loffler et al., 2021). Error propagation makes it impossible to identify relevant cellular events (such as cell death or cell division), to reliably trace lineages and to analyse signalling dynamics over long time scales. It also prevents the analysis of the correlation between shape, signalling and cell fate specification (Amat et al., 2014; Farrell et al., 2017). A few recent tools perform cell segmentation and tracking jointly, in an effort to mitigate error propagation (Bragantini et al., 2025; Ollion et al., 2024). But even these tools yield errors than can be numerous on large movies.

Concomitantly, the bioimage analysis field faces a growing need for curated datasets to develop, train and evaluate new segmentation and tracking algorithms, driven by the advent of deep learning (Maska et al., 2023). When used as ground-truth, very high quality datasets are essential to evaluate and quantitatively compare algorithm performance. When used to train a novel deep-learning model, the quality of these curated datasets will impact the training and performance of the model. However, their manual curation requires a huge amount of time and painful and unrewarding work (mostly to identify and locate segmentation errors and then correct them), which is currently the main reason behind the limited amount of available curated datasets. In addition, curation must be performed by biology experts, often experimentalists, who may not have strong computational expertise. Altogether, this calls for user-friendly tools to ease and accelerate the manual correction of such movies.

Despite this clear need, very few tools are dedicated to manual annotations and segmentation curation of epithelial movies. TissueAnalyser offers segmentation, tracking, correction and analysis options within the same framework, initially in Fiji (Aigouy et al., 2016) then transposed to Python (Aigouy and Prud’homme, 2022), but is no longer actively developed (archived repository on github, (Aigouy, 2016)). TissueAnalyser offers error correction such as junction removal or cell merging but is not well suited for large movies. Importantly, it does not interoperate with external segmentation pipelines, which limits its wide applicability. Epitools (Heller et al., 2016) ships many features to ease manual correction. It is distributed in two parts, one based on MATLAB and one on Icy (de Chaumont et al., 2012). Its use for curation requires mastering two interfaces and constant export/import of data between the two frameworks, which slows down the work. Similarly, SEGGA (Farrell et al., 2017) proposes features to segment, track, curate and analyse epithelia within one framework. However, it is not maintained anymore, and has no interoperability with other tools. MorphoGraphX (Barbier de Reuille et al., 2015) is dedicated to the segmentation and analysis of 3D epithelia and proposes a user interface and assisted correction options. However, it is developed for Linux system only, an operating system rarely used by biologists. Given these possibilities and limitations, curations of most epithelia movies are currently conducted with customed pipelines and patchwork solutions, requiring user time and pain to juggle between various software tools and perform the curation (Paci et al., 2025; Rozbicki et al., 2015), or do not offer the flexibility to integrate custom segmentation solutions as inputs and connect easily with various postprocessing pipelines.

Here, we present a new framework to ease manual curation of segmented epithelia movies called EpiCure (Epithelial Curation), guided by an epicurean philosophy of minimization of pains (O’Keefe, 2009). Its implementation is guided by direct test and feedback from several users, prompting its ergonomics and the development of specific features motivated by concrete use cases. It aims at being a versatile tool that can be interfaced with many other frameworks: it takes as input segmentations from the most classic tools and recent deep learning based softwares, and includes export options to several formats, compatible with Python and Fiji (Schindelin et al., 2012) ecosystems. In the following, we describe the general design of EpiCure, demonstrate the importance of using curated datasets for investigations in development biology, and show how EpiCure addresses the challenges and tedium of curation.

## Results and discussion

### EpiCure design

EpiCure is implemented as a Napari plugin (Sofroniew et al., 2019) to take advantage of its graphical interface, visualization capacity and interactivity features, and to connect it to recent and future segmentation tools, mostly developed in Python (**Figure 1**). EpiCure focuses on a specific image type (2D time lapses of adherent cells) but is made to be used with a wide range of biological applications. It can read data from many file formats and can integrate inputs from many external tools, allowing the use of tailored and optimized segmentation solutions. Segmentation and tracking results from state-of-the-art tools such as Cellpose (Stringer et al., 2021), EpySeg (Aigouy et al., 2020), TrackMate (Ershov et al., 2022), Trackastra (Gallusser and Weigert, 2025), Dist2Net (Ollion et al., 2024), or coming from common tools used in bioimage analysis (Fiji, napari, Python, Jupyter), can be easily loaded into EpiCure (**Figure 1A, B**). To avoid back-and-forth export / import between several external tools, we also integrated tools to perform both segmentation and tracking within EpiCure. We implemented direct connection to EpySeg (Aigouy et al., 2020) for segmentation, through a dedicated plugin, napari-epyseg that can be called within EpiCure (Letort, 2025). We also added the LapTrack algorithms for tracking (Fukai and Kawaguchi, 2023) and our own implementation of standard methods such as seed-based segmentation for local corrections (**Figure 1B**).

**Figure 1:**
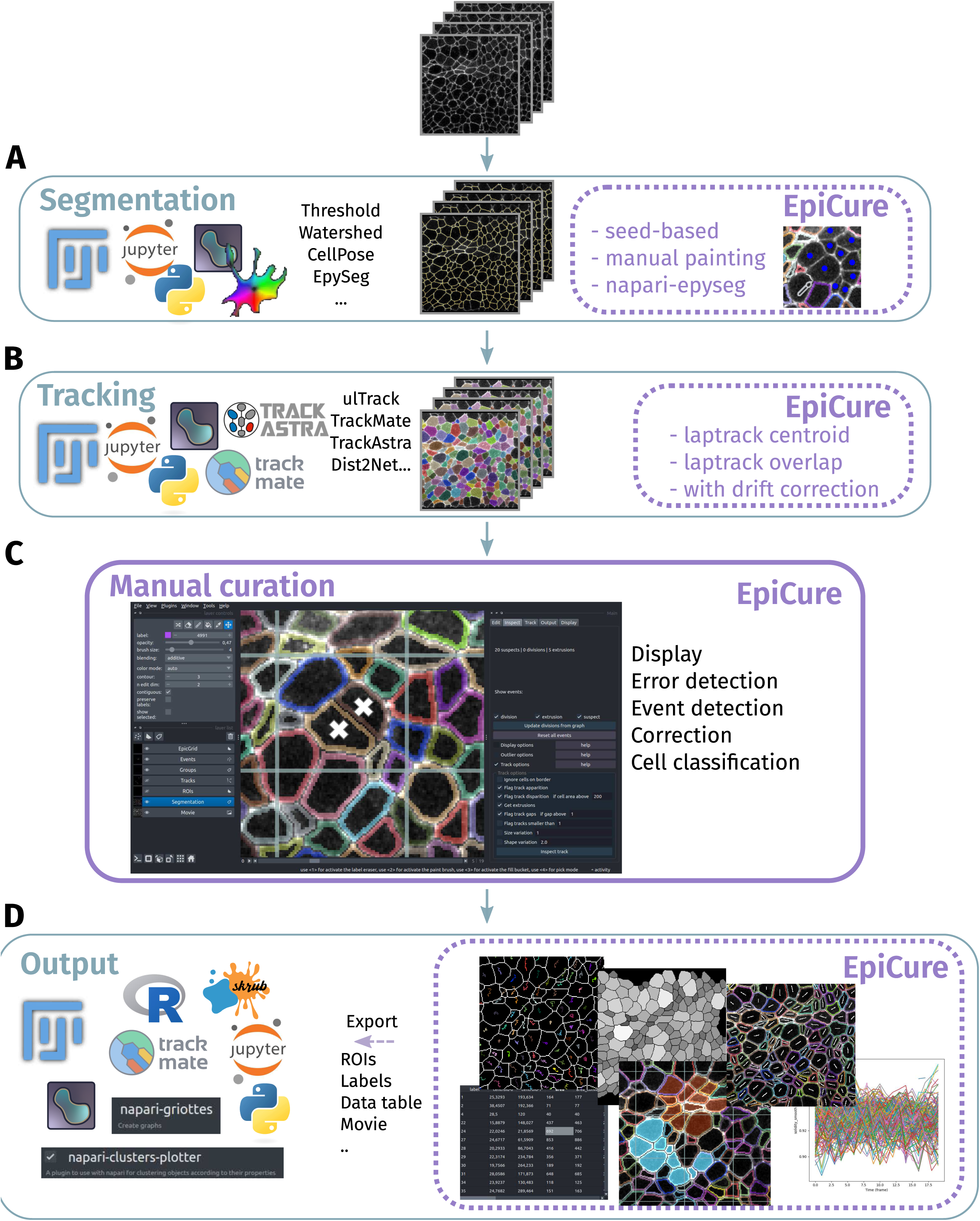
EpiCure’s overview. (A) Segmentation step. Segmentation from a variety of external softwares can be imported or segmentation can be performed within EpiCure through several options. (B) Tracking step. Tracking results from a variety of external softwares can be imported or tracking can be done within EpiCure based on the Laptrack module. (C) Main core of EpiCure: visualisation and curation of the segmentation and tracking. EpiCure interface as a napari plugin. The movie and the different layers are overlaid in the central part of the window. On the left side, the Napari panel displays the layers (raw movie, segmentation, tracks..) that can be set visible or not and for the selected layer, The Napari options are available at the top. On the right side, the EpiCure panel proposes 5 interactive tabs to display, inspect, edit, track and exploit the results. (D) Output step. After correction, the segmentation and tracking can be directly measured and plotted within EpiCure or exported to other external softwares to perform further analyses.

The central goal of EpiCure is to ease manual correction of epithelia segmentation of cells tracked over time, which we achieve with a handy user interface. EpiCure can be controlled via the graphical interface and / or with mouse and keyboard shortcuts for fast and convenient usage. To keep the data overlay clear and lean, the main display has several modes that correspond to different steps in curation (“Display” features, **Figure 1C, Movie S1**). Alternating between these modes, the user can detect, show and navigate possible segmentation/tracking errors (“Error detection”), detect / correct cellular events as cell division or extrusion (“Event detection”), perform segmentation correction with one or two clicks (“Correction”) and group cells by user-defined type or automatic classification (“Cell classification”).

EpiCure also enables analysis of the curated data, either externally through napari, Fiji, or analysing the results in Excel, Python and R, or internally in EpiCure. EpiCure proposes several export formats compatible with many softwares that can exploit its data (**Figure 1D**). Common analyses such as measuring cell morphological properties, tracking properties, measuring intensities and plotting their evolution over time can be performed directly within EpiCure (**Figure 1D**), avoiding the need to export and switch to another tool for previsualisation. Specific analyses are additionally proposed as Jupyter notebooks directly interacting with EpiCure (**Supplemental Information).** These notebooks can easily be adapted to address specific analyses. Overall, while the core of EpiCure resides in its ergonomics to efficiently help curation of segmentation and tracking errors, it amounts to a fully featured tool, interoperable with the key softwares in bioimage analysis, and allows performing the whole analysis that goes from raw images to single cell measurements and dynamic plots.

In the following, we will describe EpiCure’s main features in greater detail, illustrated with use-cases for each feature and discussing the importance of manual corrections for downstream analysis.

### EpiCure highlights possible segmentation errors for faster correction

A typical movie of epithelial development can contain thousands to millions of “objects” (individual cells), which have to be followed over time (**Figure 2A**). Tracks can be directly loaded into EpiCure, or cells can be directly tracked within our plugin through the “Track” panel, based on the LapTrack library (Fukai and Kawaguchi, 2023). Several LapTrack algorithms can detect divisions during tracking and allow adding constrains such as linking cells only with a similar area / shape. EpiCure associates with each cell and track a unique ID and unique colour in the interface (through the Label layer of napari). Cells alone can be displayed or overlaid with the associated track as a line showing the local movement of the cell (**Figure 2A**). EpiCure also offers several functionalities to switch rapidly between display modes (full cell coloured, only cell contour, with / without tracks, visible division events (**Movie S1**) offering flexibility for the visual inspection of the results.

**Figure 2:**
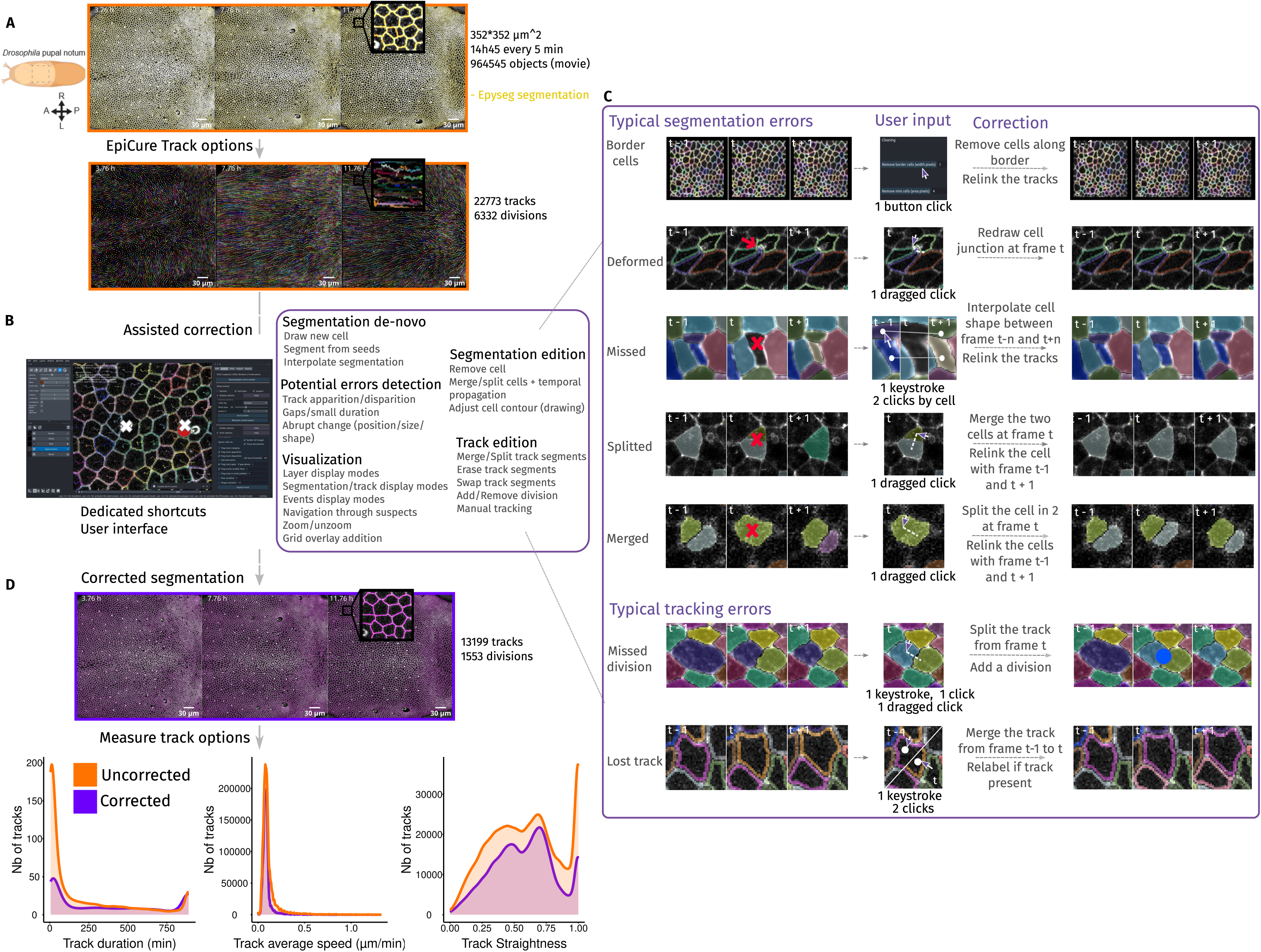
Assisted correction using temporal information. (A) Top: Snapshot of local z-projection of a *Drosophila* pupal notum movie from 20 to 32 hours after pupal formation, cell contours are marked by E-Cad::GFP (grey). The segmentation obtained with Epyseg is shown in yellow. Inset shows a closest view with a segmentation error in the middle (additional junction). Bottom: segmented cells are tracked and tracks are displayed as colored lines within EpiCure. Scale bar=30μm. (B) Assisted correction features using the temporal information from tracking. Overview of EpiCure interface and list of the main features to ease manual correction of the segmentation, sorted into 5 categories: segmentation de-novo, segmentation edition, potential error detection, visualisation and track edition. (C) Examples of the main typical segmentation (top) or tracking (bottom) errors. For each typical error (one line), the user input required to correct the error easily is described by the number of clicks or keystrokes (column “User input”). Automatic treatment (column “Correction”) is applied following the user’s input resulting in the corrected segmentation and tracking. (D) Results of the segmentation curation. Top: raw movie (gray) overlaid with the curated segmentation (magenta), inset shows the same close-up view as in (A). Bottom: Comparison of quantitative measurements with EpiCure before (orange) and after (purple) curation: histogram of track duration (left), average speed of the tracks (middle) and track straightness (net displacement/total displacement, right). Scale bar=30μm.

Finally, EpiCure takes advantage of the temporal information to detect potential errors of segmentation. Tracking is more likely to fail where cells were wrongly detected (Bragantini et al., 2025; Loffler et al., 2021). Inspecting the tracks for unexpected disappearance / appearance or sudden change of cell property (Loffler et al., 2021) can point to potential errors. To ensure flexibility, users can configure in the interface which suspicious track changes should be flagged (**Figure 2B**). The user can then scroll the whole movie and zoom in and out on potential errors or directly use dedicated keyboard shortcuts to navigate through all the potential errors, hence fastening error detection and correction.

### EpiCure assists and propagates segmentation correction

Several options are proposed to rapidly fix true segmentation errors (**Figure 2C**). For instance, cells are regularly split into two cells in a single frame due to over-segmentation (“Split”). By simply doing a dragged click from one cell to the other, EpiCure will merge them into a single cell. It will then directly try to relink the corrected cell with the previous and following tracks, propagating the local correction to the tracking information. The most typical segmentation errors as well as tracking errors are all covered by the plugin with features for correction involving only one or two keystrokes or a simple mouse click, while automatically relinking the tracks after correction (**Figure 2C**). Of note, these typical errors and correction features rely on *in vivo* epithelia properties: namely that cells are contiguous and connected by apical junctions. This geometric constraint allows to reduce considerably the correction efforts, by proposing options to draw only the junctions without having to redraw the cells entirely. It should be noted that as a result, EpiCure is not optimised for the correction of non-confluent cells.

To illustrate the importance of manual segmentation correction of cell tracking, we analysed some track features before and after correction in a movie of the *Drosophila* pupal notum (a single layer epithelium) segmented with EPySeg (Cumming et al., 2025) (**Movie S2**, **Figure 2D**). Curation of segmentation drastically changes the number of tracks and track duration, yielding much longer tracks (**Figure 2D**) (Farrell et al., 2017). Some temporal measurements are not highly sensitive to fragmented tracks, such as the average cell speed, and are therefore mostly unaffected by segmentation and track correction (**Figure 2D**). Other features, however, require accurate tracking of cells at long term, such as lineage tracing, cell divisions and cell deaths localisation, or measuring temporal change of shape and signalling. For them, the correction of errors becomes essential and drastically changes the analysis output. Segmentation error correction allows to extract accurate quantitative information about tracks as well as the variation of cellular features over time (morphology, signal intensity variation, see below).

### The impact of segmentation error on rare cell types

EpiCure includes options to measure and plot features of tracked cells directly in the interface (**Figure 3A**), focusing on features relevant for epithelia cells, such as the number of neighbours (Farhadifar et al., 2007), the shape index (Bi et al., 2016), junctions or the cytoplasmic intensity of relevant markers. All these measures are displayed in interactive tables. For user convenience, each feature can be displayed as a coloured temporal map in which each cell is coloured by its feature value (**Figure 3B**), allowing rapid preliminary exploration of the spatiotemporal distribution of a given feature.

**Figure 3:**
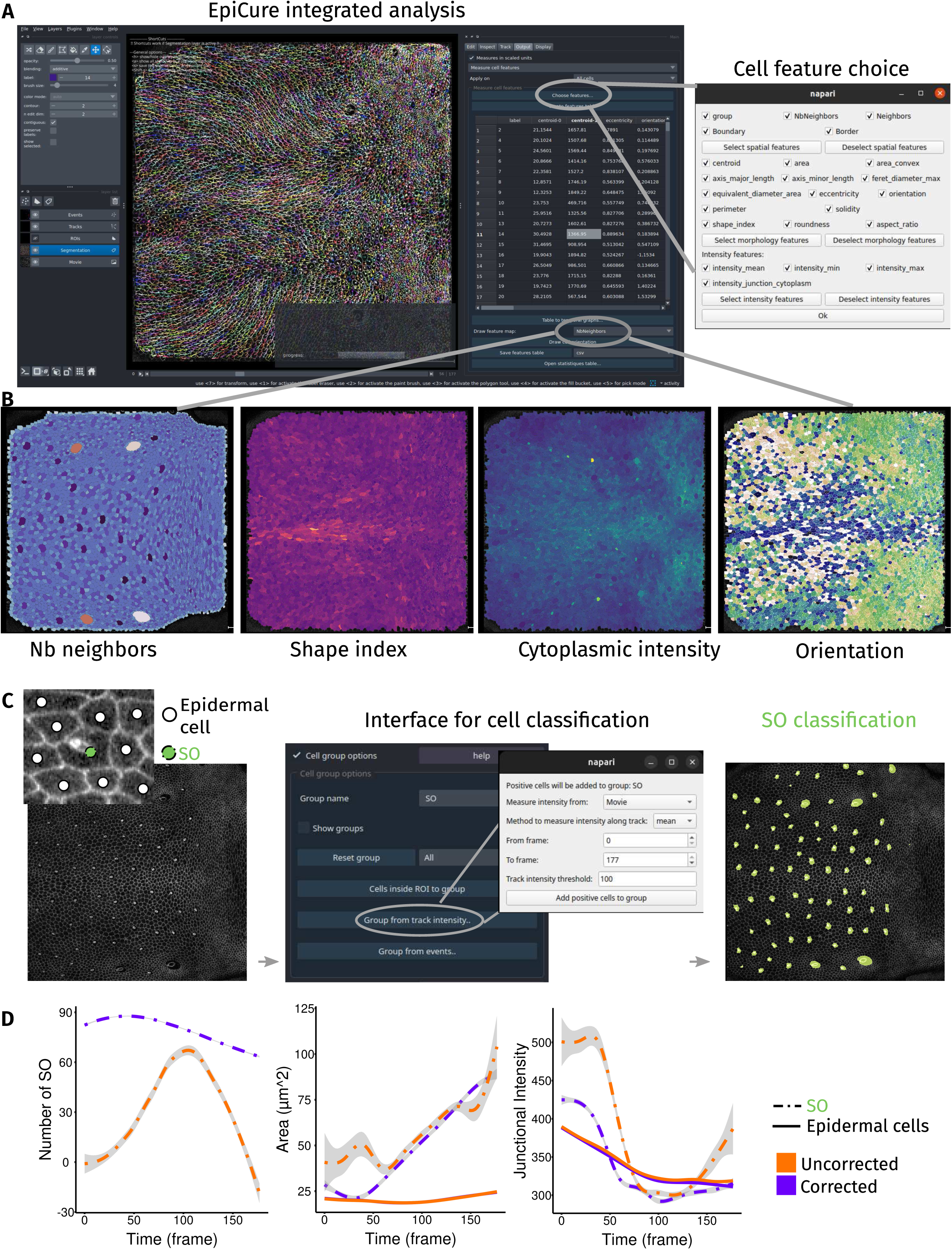
Impact of corrections on cell scale quantifications. (A) Interface for cell scale measurement. Left: snapshot of EpiCure interface with visible coloured tracks and the measurement option panel at the left of the window. Right: panel that pops-up when “Choose features” button is clicked, to choose which cell characteristic to measure. (B) Examples of measurement heatmaps. Each cell is coloured by the relative value of the selected feature: number of neighbours of the cell (left), shape index (middle left), cytoplasmic intensity (middle right) and cell main orientation (right). Different colourmaps have been chosen for each heatmap. (C) Cell classification as SO cells (Sensory Organ cells) or epidermal cells. Left: snapshot of the whole notum at 32 hours after pupal formation. Left top panel: example of one SO (light green dot) surrounded by several epidermal cells (white dots). Middle: EpiCure group panel to manually/automatically define cell classification into groups. Clicking on “Group from track intensity” button pops-up an interface (right) to choose parameters to automatically classify cells into a given group (here SO) based on intensity (here mean intensity along track, above a threshold). Right: Results of the classification with SO cells colored in light green. (D) Comparison of temporal quantifications before and after correction. Temporal evolution of cell scale features measured with EpiCure before (orange) and after (purple) curation: number of objects classified as SO (left), average cell area (middle, encompassing all the cells of the SO) and average junctional intensity (average of each cell intensity along the junctions, using the most external cell, the socket, for SOs, right). Dashed lines represent the measure of SOs only and full lines the measure of all other epidermal cells. Grey area is the confidence interval at 0.95

Analysing developing tissues often requires extracting parameters from specific cell populations, such as distinct cell types or fates, or clonal perturbed cells in mosaic tissues. To facilitate such analyses, we added an option to classify cells into user-defined groups. Cells can be automatically classified based on a marker’s intensity (**Figure 3C**), on their fate (e.g., cells that will divide or extrude) or manually through a keyboard shortcut. To illustrate this option, we automatically (see **Material and Methods**) identified the apical area of Sensory Organs (SO, precursors of sensillum generated by successive asymmetric divisions (Hartenstein and Posakony, 1989)) in the *Drosophila* pupal notum in comparison with normal epidermal cells (**Figure 3C**). We measured the cell morphologies over time and compared their value for cells marked as SOs (**Figure 3D**, dashed lines, encompassing the full apical area of all the SOs cells) vs other epidermal cells (**Figure 3D**, full lines). To illustrate again the importance of correction for quantitative features extraction, we repeated this analysis on the segmentation before (**Figure 3D**, orange lines) and after correction (**Figure 3D**, magenta lines). At the beginning of the movie, many SO precursors were not recognised without correction (**Figure 3D**, left panel), because SOs are only morphologically recognisable at late development (Besson et al., 2015) and the tracks were lost before SO could be identified. Moreover, toward the end of the movie, the socket cells encompassing the rest of the sensory organ have a much larger apical area compared to other cells (**Figure 3D**, middle panel) and are quite systematically over segmented. In uncorrected movies, the low SO cells detection rate coupled with high SO segmentation error rates strongly affects the quality of feature measurements such as full apical area of the SO or junctional intensity of the socket cell (**Figure 3D**, dashed lines), while measures of other epidermal cells were much less impacted by segmentation errors (**Figure 3D**, full lines). This boils down to SO cells being more error-prone in the segmentation due to their atypical shape and under-representation in the segmentation training data. This use-case highlights the importance of curating atypical groups of cells (*e.g.* clonally perturbed cells, differentiated cells) that are more likely to be wrongly segmented by deep-learning pipelines, and the challenge of segmenting heterogeneous tissues.

### Curating dataset also helps recognise cellular events

Understanding morphogenesis requires precise quantification of the spatio-temporal patterns of cellular events such as cell division and extrusion/death (Etournay et al., 2016; Guirao et al., 2015). While inspecting tracks for possible segmentation errors, extrusion or division events can also be detected and displayed in EpiCure (**Figure 4A**). Interestingly, cell division and cell extrusion are also quite frequently a source of segmentation or tracking errors (**Figure Sup 4A**) and are also most of the time wrongly estimated when quantified based on uncorrected tracks (Amat et al., 2014; Schiegg et al., 2013). Manual inspection of potential segmentation errors allows detecting events that could be missed by the automatic detection of cell division or cell death from track analysis. Therefore, we integrated user-oriented features in EpiCure to be able to display, navigate and edit these single-time-point events (**Figure 4A**). We compared their spatial distribution when their detection solely relies on automated analysis versus after manual curation (**Figure 4B, Movie S3**). This comparison shows that extrusion events are accurately captured without correction (**Figure 4B,C, Movie S3**). In contrast, a high number of false positive events for cell divisions are detected, mostly caused by erroneous tracking (**Figure 4B,C, Movie S3**). Similarly, we explored how the relative cell apical area of the central cell evolved relative to its neighbours until cell division/cell extrusion termination (**Figure Sup 4B**). While this measurement was again not much affected by curation for extruding cells, the strong increase of cell size just before division was not properly captured without segmentation curation (most likely because many non-dividing cells where incorrectly classified as dividing and incorporated in this measurement). Together with the previous observation on SO morphology, these results demonstrate that manual correction is an important step as soon as the analysed features (cell type, morphology, events) are sensitive to possible segmentation errors.

**Figure 4:**
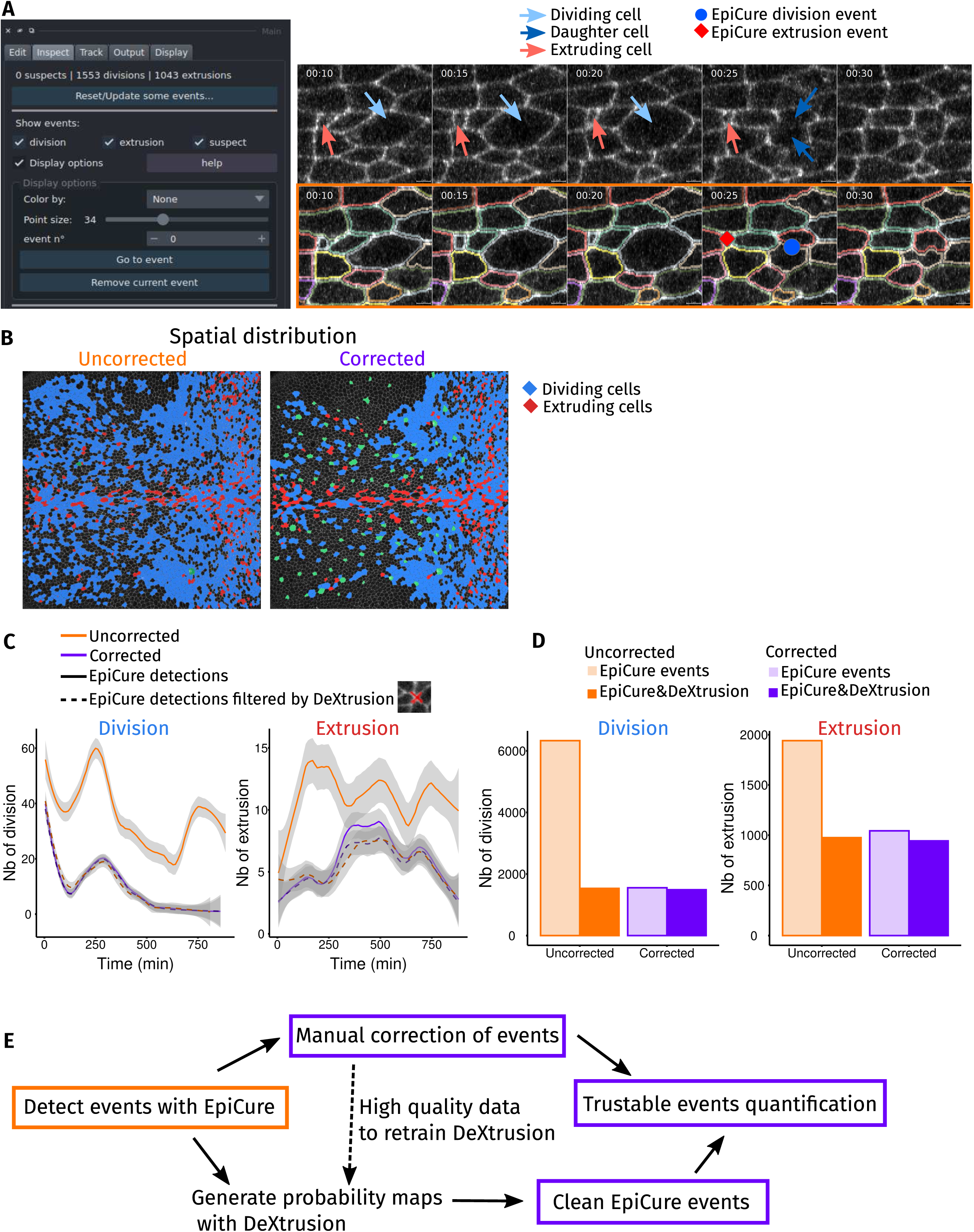
Quantification of cellular events (divisions, extrusions) (A) Visualization of cellular events. Left: interface to choose the cellular events display and navigate through them. Right: example of cell division (blue arrows and dot) and extrusion (red arrows and dot) on the raw movie (top) and the segmented cells (bottom). Division events are placed in the middle of the two daughter cells at the first time of their appearance. Extrusion events are placed in the extruding cell at the last frame where the cell apical area is still visible. (B) Impact of segmentation curation on events spatial distribution. Spatial distribution of cells that will divide later (blue) or extrude (red) on the first frame of the movie, for uncorrected analysis (left, orange) and after segmentation curation (right, magenta). The Sensory Organs (SOs) are shown in green. (C) Temporal distribution and interoperability with DeXtrusion (Villars et al., 2023) for detections filtering. Temporal evolution of the number of detected divisions (left) and extrusions (right) on uncorrected (orange) or corrected (purple) movies. Events are detected with EpiCure (full lines) or also filtered for events having also a high probability prediction in DeXtrusion (dashed lines, events detected by both pipelines). (D) Interoperability with DeXtrusion for detections filtering. Total number of divisions (left) or extrusions (right) for the uncorrected (orange) and corrected (purple) movies. Events are detected with EpiCure (light colors) or filtered with DeXtrusion (dark colors). (E) Pipeline for a reliable events quantification. Steps to have reliable events quantification: detecting events with EpiCure and manually correct them, the curated events can then be analysed. This can also serve as a training database to fine-tune DeXtrusion to new datasets. Another possibility to avoid manual correction is to generate probability maps with DeXtrusion and use them to filter events that are detected by both softwares. This directly generates a set of clean cellular events that can be used to track the dynamics of cells undergoing those events.

Since the identification of extrusion events was reliable despite segmentation errors, while divisions were largely overestimated, we investigated whether we could filter these events in EpiCure to keep only “probable” events without manual correction. We relied on DeXtrusion (Villars et al., 2023), a deep-learning pipeline that we previously developed to detect cellular events on raw movies, without segmentation and tracking. DeXtrusion provides probability maps of the presence of division and extrusion events. We therefore used DeXtrusion probability map to filter events detected in EpiCure (before and after segmentation correction), only keeping the events also associated with high probability values in DeXtrusion. This filtering was very efficient in removing all false positive cell divisions, despite the absence of segmentation curation (**Figure 4C,D**, dashed lines and full colours): the number of detected divisions and extrusions were nearly the same for both uncorrected and corrected segmentation when cross-filtered with DeXtrusion (**Figure 4C,D**, dashed lines and full colours). This illustrates the interoperability of EpiCure, whose outputs can be cross-compared with deep learning-based detection of events, allowing to filter these events and generate quantifiable datasets without heavy manual curation (**Figure 4E**). This also demonstrates the usefulness of EpiCure to generate high quality data for retraining deep-learning tools such as DeXtrusion. Note that we did not integrate DeXtrusion directly within EpiCure as we aimed to keep EpiCure general. However, interactions between EpiCure and DeXtrusion are easily managed through available Jupyter notebooks (**see Material and Methods**).

### EpiCure: a general-purpose tool

As mentioned above, we aimed at developing a general-purpose tool for epithelial tissue segmentation curation, convenient for users working with different biological samples, imaging conditions and analysis objectives. Accordingly, EpiCure was developed through the feedback of users working with various biological systems and imaging conditions, encompassing the *Drosophila* pupal notum (Valon et al., 2021) (**Movie S2** and **S3, Figure Sup 5A**), the quail gastrula extraembryonic tissue with a membrane marker (Michaut et al., 2025) (**Movie S4**), *Drosophila* abdominal histoblasts marked by E-cad (Phan et al., 2024) (**Movie S5**) and the zebrafish telencephalic germinal tissue marked by Zona occludens 1 (Zo1) (Mancini et al., 2023) (**Movie S6**). We asked EpiCure users to measure the duration of their manual corrections using EpiCure on small, cropped images from their biological systems (**Figure 5, Figure Sup 5A, Movies S4, S5, S6**). EpiCure was run on the three main operating systems for these tests (MacOS, Ubuntu and Windows) and with both Python 3.9 and 3.10 versions (**Figure 5**), demonstrating its deployability. Each image was first segmented and tracked with different algorithms (*e.g.* Cellpose) or with EpiCure integrated tools (EpySeg), illustrating again the flexibility of the pipeline inputs. The times needed for correction were very diverse (from minutes to hours) and influenced by the diversity of input movies properties (quality of the segmentation, signal to noise, spatiotemporal resolution, morphological and dynamical properties of the biological tissues, **Figure 5** and **Figure Sup 5B**) and the users proficiency with EpiCure. Of note, the long time required to curate the telencephalon movie probably boils down to its very low temporal resolution (1 frame / 60 hours) making the association of cells between time point very challenging. To facilitate EpiCure usage, we also included an option to personalise keyboard shortcuts so that each user can optimize the combination of correction shortcuts and keyboard position. The full list of shortcuts can be opened in a separated window and is detailed for each step of EpiCure in the online documentation.

**Figure 5:**
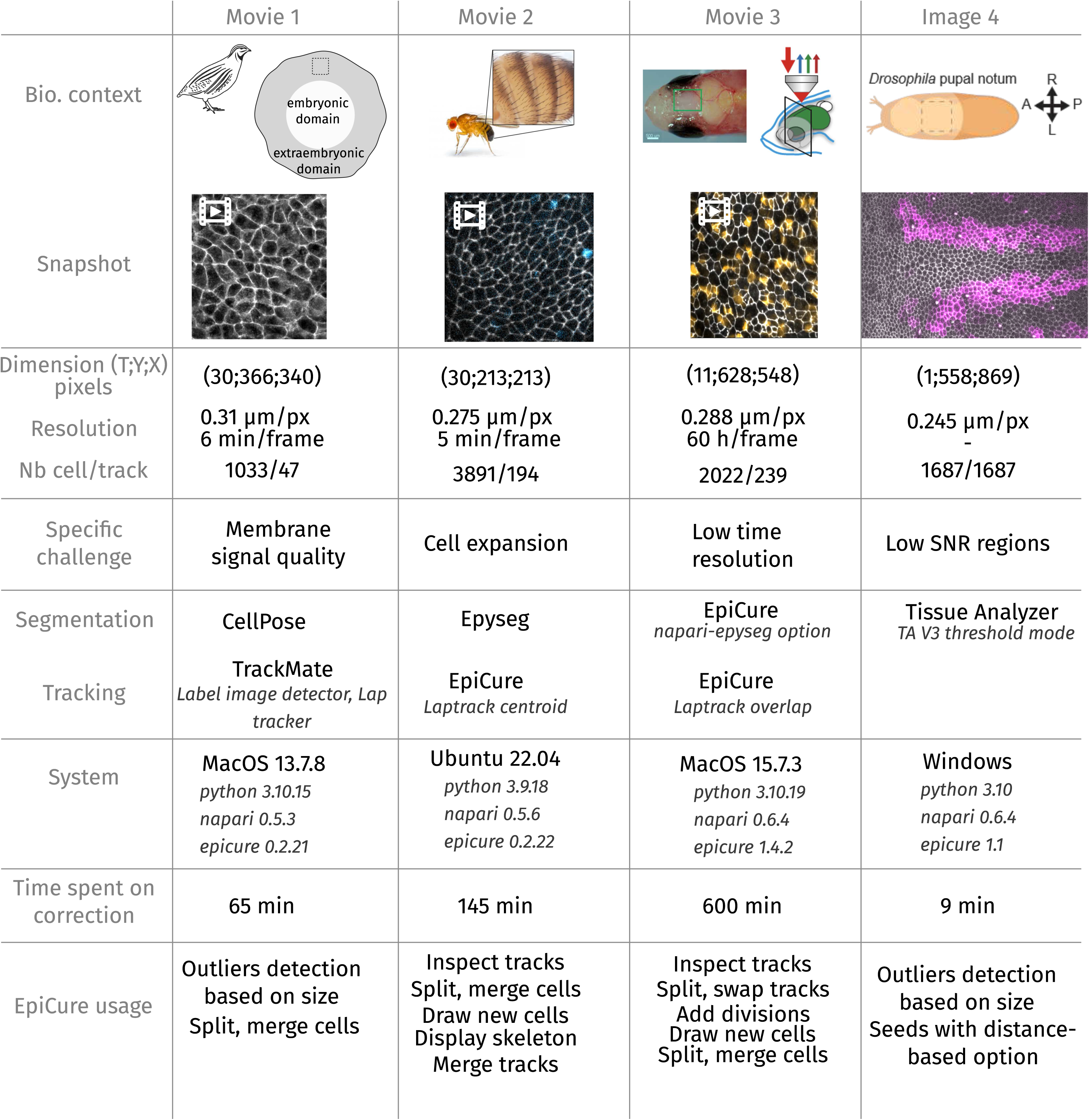
Generalisation of the usage of EpiCure. Comparison of EpiCure performance on 4 model systems, including movies of the quail gastrula marked with membrane GFP (grey, left), the *Drosophila* abdominal histoblasts marked with E-cad::GFP (grey, left, middle), the zebrafish adult telencephalic stem cell layer marked with gfap:Zo1::mKate2 and deltaA::eGFP (grey and orange, right, middle), and a single snapshot of the *Drosophila* pupal notum with cells expressing RFP in the stripe-Gal4 domain (magenta) and E-cad::GFP (grey) (right). For each system, the dimension in time frames and pixel size is provided, the resolution, the frame rate and the number of objects segmented (cells multiplied by the number of frames). We listed the specific challenge associated with each example. The softwares used for segmentation inputs prior to EpiCure are listed, as well as the tracking methodology, the operating system and Python, Napari and EpiCure versions. Each user had to estimate the total time required to perform the full curation of their segmentation and list the functionalities they mostly used in EpiCure. Note the large time variation depending on the movie/image quality and user training with EpiCure.

Quantitative single-cell studies in developmental biology require a level of segmentation and tracking accuracy that automated bioimage analysis tools cannot achieve alone. We bridge this need with EpiCure, a napari plugin dedicated to the manual curation of segmentation and tracking results, within an interface designed to accelerate the curation process and make it as streamlined as possible. EpiCure has an interactive and ergonomic interface designed for a wide audience. It was tested and adopted by several users working on many biological models, compatible with the main operating systems and several Python versions used in research. The code is open source and available on Github (https://github.com/Image-Analysis-Hub/Epicure/), with dedicated online documentation for installation and usage (https://image-analysis-hub.github.io/Epicure/). Additionally, the plugin provides in-interface help windows for every function of the graphical user interface, facilitating its quick adoption. Keyboard or mouse shortcuts are proposed to ease the process and can be customised by the user. The plugin is compatible with a wide range of inputs (**Figure 1**, **Figure 5**) and can export segmentation, tracking and analysis results to various formats (**Figure 1**), making the pipeline both self-sufficient and interoperable with other existing and future softwares.

One potential issue for EpiCure wide adoption might rely on its installation, which is not always straight-forward for users who are not familiar with Python. We therefore created clear installation instructions and paid particular attention to limiting EpiCure dependencies to reduce the risk of incompatibilities. The most potentially problematic dependency, EpySeg (limited to python 3.10 and relying on tensorflow) is handled automatically by EpiCure in a dedicated virtual environment thanks to the appose library (https://github.com/apposed), making its installation invisible for the user. Napari can now be installed as a downloadable, installable bundle app for all the major OSes used in research, and from which EpiCure can be installed seamlessly. Another limitation of EpiCure stems from its focus on 2D epithelial tissue movies and confluent cells. We also integrated one option allowing the user to run it on non-confluent cells, but many curation features of EpiCure are not easily amenable to such use-cases. Bridging them within EpiCure is an interesting endeavour for future versions. Our primary objective was to develop a versatile, general-purpose tool for correcting epithelial segmentation and tracking, targeting a widespread bottleneck encountered by multiple research groups studying 2D epithelia, such as those in quail, *Drosophila*, or zebrafish during development and beyond (**Figure 5**). Indeed, these datasets represent some of the largest and most computationally demanding time-lapse sequences currently generated in developmental biology. We therefore decided to avoid relying on a purely automated deep-learning solution, which requires training on representative data for each type of sample and imaging modality, and computational resources not necessarily accessible to every user. However, deep-learning pipelines do show interesting performance for automatic error correction or assisting manual annotations (Betjes et al., 2025; Isensee et al., 2025; van der Wal et al., 2021). Future versions of EpiCure could integrate such solutions as optional features, but they will require retraining the deep-learning models for each type of dataset. Finally, EpiCure was designed as an interoperable component that can easily be connected with other existing and novel pipelines while facilitating adoption of the code by developers who may be willing to add new options and integration in future versions. Developer-oriented documentation (API) is available on the documentation pages (https://image-analysis-hub.github.io/Epicure/api/epicure.html) to facilitate contributions. This will ensure constant evolvability and maintenance of the tool and long-term relevance for the fast-evolving field of tissue segmentation.

## Acknowledgements

Work in JG lab is supported by the Institut Pasteur, the ANR ForcePattern, the ANR CellTeam the ANR-10-LABX-0073, the CNRS (MR3738) and the European Union (ERC MechanoselfFate). AM was supported by a Pasteur-Roux-Cantarini postdoctoral fellowship. Work in FS lab was supported by the ANR-10-LABX-0073, the CNRS and the Institut Pasteur. LX received funding by the Fondation pour la Recherche Médicale (FDT202504020138) and by the INCEPTION project (PIA/ANR-16-CONV-0005), and is a student from the FIRE PhD program funded by the Bettencourt Schueller foundation and the EURIP graduate program (ANR-17-EURE-0012). TC was supported by a grant from the PPU program of Institut Pasteur and a 4^th^ year PhD grant from the Ligue Nationale Contre le Cancer. Work in RL lab is supported by the Institut Pasteur, the ANR PRC CoECECa, the ANR PRC MAPEFLU, the ANR-10-LABX-0073, the CNRS (UMR 3738) and the European Union (ERC, PrApEDoC, 101085444). Work in LBC’s lab is supported by Institut Pasteur, CNRS, the Agence Nationale de la Recherche (ANR-10-LABX-0073), the Fondation pour la Recherche Médicale (EQU202203014636), the European Research Council (ERC SyG PEPS 101071786). Work of CTR is supported by the Chan Zuckerberg Initiative (CTR) and by National Institutes of Health Grant P41GM135019. We acknowledge France-BioImaging infrastructure (https://ror.org/01y7vt929) supported by the French National Research Agency (ANR-24-INBS-0005 FBI BIOGEN). Views and opinions expressed are however those of the author(s) only and do not necessarily reflect those of the European Union or the European Research Council. Neither the European Union nor the granting authority can be held responsible for them. A CC-BY public copyright license has been applied by the authors to the present document and will be applied to all subsequent versions up to the Author Accepted Manuscript arising from this submission, in accordance with the grant’s open access conditions.

## Authors contributions

GL and RL designed the project and coordinated its execution. GL performed all the code development, user interactions and pipelines integration. TC and LV provided segmented data for the pupal notum. AM and JG provided segmented data for the quail. ND and LBC provided segmented data for zebrafish. MSP and FS provided segmented data for the *Drosophila* pupal abdomen. TC, LV, AM, MSP and ND provided regular feedback to adapt the pipelines. LX developed the code for import and export between EpiCure and TrackMate. CTR developed the appose library allowing seamless execution of EpySeg irrespective of the Python version. JYT and LBC helped to monitor the project and define the positioning strategy. All the authors participated in the article writing and editing.

## Material and Methods

### Biological dataset

Movie of the pupal notum was taken from the following reference where experimental details can be found (Cumming et al., 2025). Movie from the gastrulating quail was taken from (Michaut et al., 2025). Movie from the *Drosophila* abdomen histoblast was taken from (Phan et al., 2024). Movie from the zebrafish telencephalon was taken from (Mancini et al., 2023).

### Tool implementation and availability

EpiCure is a Python module developed as a napari plugin (Sofroniew et al., 2019). It can be installed via pip or in the napari plugin interface. It is distributed open source under the BSD-3 License and can be downloaded here (https://github.com/Image-Analysis-Hub/Epicure/). Instructions for installation and detailed usage are given in the online user documentation (https://image-analysis-hub.github.io/Epicure/).

### Analysis

Quantitative measures presented in this paper were extracted from EpiCure Output’s option (https://image-analysis-hub.github.io/Epicure/Output/) and plotted using the R software.

#### Track feature analysis (Figure 2)

data was exported with EpiCure “measure track features” option as a .csv data file. From this file, we extracted and plotted the histogram of the track duration (total duration of one track from its appearance to the cell division/extrusion/disappearance from the movie), average speed (cell speed of motion between two consecutives frames, averaged over the whole track) and straightness (distance between the last and the first position of the track, divided by the total distance travelled by the cell).

#### Sensory Organ (SO) analysis (Figure 3C-D)

to classify cells as SOs, we took advantage of our recently developed pipeline, DeXtrusion (Villars et al., 2023), which is dedicated to the detection of cellular events such as cell extrusion and can in particular generate probability maps of SO presence. We loaded DeXtrusion probability maps of SO presence in EpiCure and used the “Group from track intensity” feature of EpiCure (in “Edit” panel) to classify cells that had a high probability along the entire track of being a SO. A few (around 5) missed cells were manually added to the group (**Figure 3C**). This interaction DeXtrusion/EpiCure was done through a Jupyter notebook available in the github repository (https://github.com/Image-Analysis-Hub/Epicure/blob/main/notebooks/EpiCure_DeXtrusion_Interaction.ipynb). Cell properties such as cell area and shape index were exported to a .csv table through the “measure cell features” option of EpiCure (in the “Output” panel), along with their group classification as SO or not. This table was then loaded in R to generate the temporal plot of the evolution of these features in time in the movie (**Figure 3D**). Temporal plots can also be generated within EpiCure (with the “Table to temporal graphs” buttons in the “Measure cell features” option).

#### Biological events analysis (Figure 4)

Divisions can be detected via the tracking algorithm, if done within EpiCure with the LapTrack option. It is also possible to find them through track inspection, in the “Inspect” panel of EpiCure with the “get divisions” option. It will detect a division when one track disappears and two very nearby tracks appear. Similarly, extrusions are marked when a track disappears without a division and with a small area as the cell apical size decrease when extruding. Cells can be classified by their fate regarding this event, with the “Group from events” option of the “Edit” panel of EpiCure. It will mark all cells that will divide in the group “Dividing” and cells that extrude in the group “Extruding” (**Figure 4B**). To measure the cell’s relative area (compared to its neighbours) from the track appearance to its fate (division or extrusion), the list of events (division and extrusion) and their spatiotemporal position was exported with the “export/measure events” option of the “Output” panel. The table of cell area and their neighbour at all time points was exported with the “measure cell features” options, with the features “area” and “NbNeighbor” selected. These two tables were put together for analysis of the cell area divided by the mean area of its neighbor cells, at all times until the event. This analysis is proposed in the notebook “PostEpiCureAnalysis_extruding.ipynb” in the github repository. The resulting time plots were then plotted in R but they can be directly generated in the notebook. Finally, the temporal evolutions of the number of events (**Figure 4C**) were plotted in R from the events table exported with the “export/measure events”. These temporal curves of the number of events can also be plotted directly within EpiCure.

#### Generality of EpiCure usage (**Figure 5, Figure Sup 5)**

To estimate typical proportion of segmentation/tracking error and the time necessary for correction with EpiCure on several dataset, 4 researchers from different labs were asked to time their manual curation of a cropped movie/image of their biological system. They reported the time necessary for the correction, their system configuration (operating system, tools version), and the segmentation and tracking tools used. The proportion of segmentation error was then estimated by comparing the initial segmentation + tracking (before correction) and the resulting segmentation + tracking after correction. Both segmentations (before/after) were exported from EpiCure as label movies that were compared. We measured the skeleton intersection-over-union (IOU) score: for each movie (before/after) correction, the skeleton of the segmentation was exported from EpiCure and the binary mask IOU was calculated. This measure is only sensitive to segmentation error. CellNbError: the total number of cell/object (over the whole movie regardless of the track) segmented in the movie before and after correction were compared as the total number of cells after correction (true number of cells) minus the number of cells before correction (initial segmentation) divided by the number of cells after correction. This measure is only sensitive to segmentation performance. LengthTrackError: the error on the measure of the track length (total duration) was estimated as the average track duration after correction (ground truth) minus the average track duration before correction (initial tracking) divided by the average track duration after correction (ground truth). This measure reflects how the segmentation and tracking errors affect quantitative measurements such as track duration. AreaTrackError: the error on measuring cell’s average area on uncorrected movie was estimated as the average cell area along tracks in the movie after correction (true average area) minus the average cell area before correction (initial segmentation) divided by the average cell area after correction. This measure reflects how the segmentation and tracking errors affect quantitative measurements such as average cell size.

---

**Figure Sup 4:**
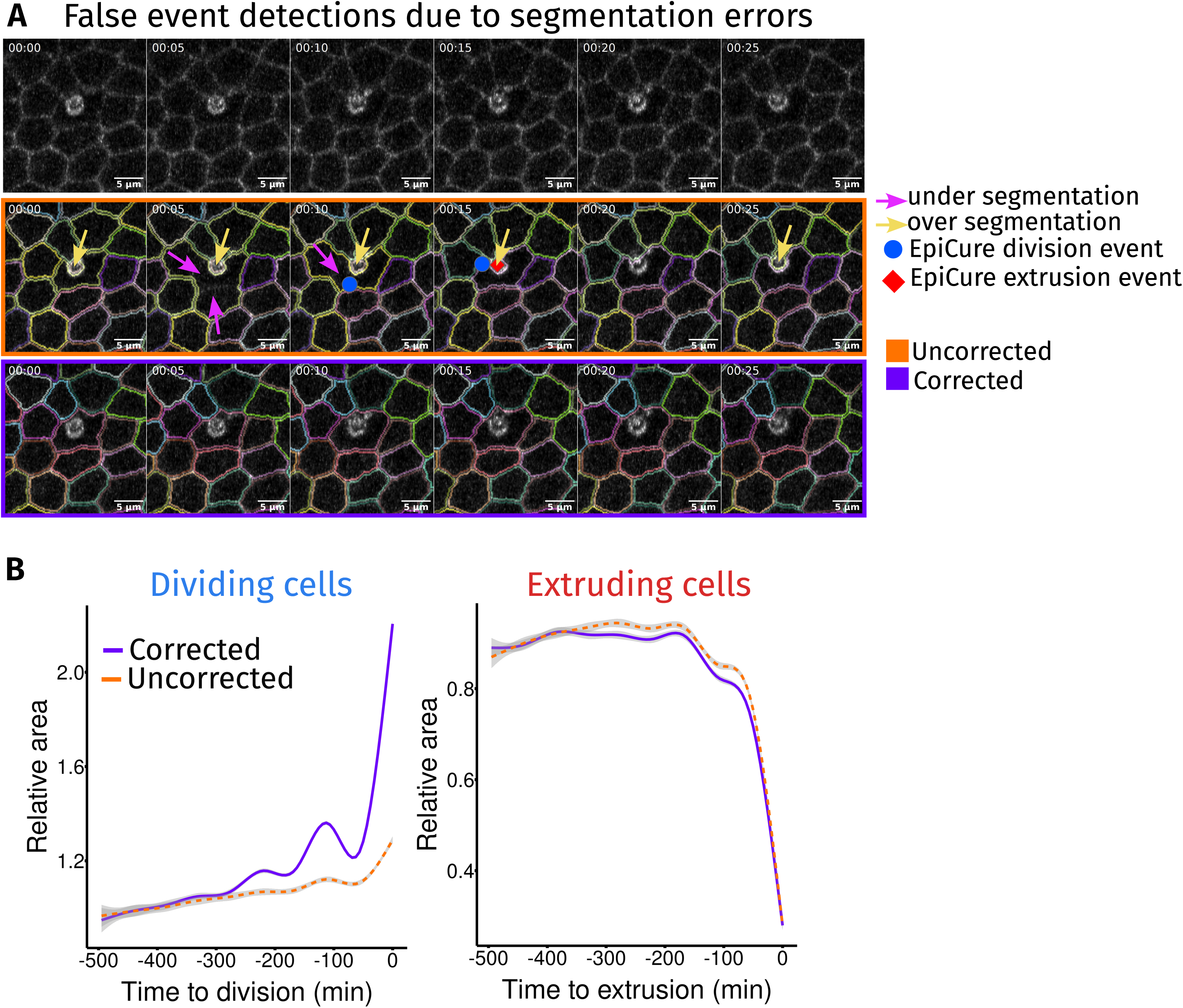
Impact of segmentation errors on cellular events detection. (A) False event detections created by segmentation errors. Top: crop of the raw movie on a zone of interest. Middle: Segmented movie without correction: under segmentation is visible in three parts of the movie, where junction signal is faint (pink arrows). Over segmentation is visible in 5 frames, due to the SO cells (yellow arrows). These errors generate wrong tracking and thus false events (division and extrusion) detection. Bottom: Corrected segmentation. No events are detected anymore in this cropped movie. (B) Impact of segmentation correction on events temporal distribution. Temporal evolution of the cell area relative to the area of its neighbours, for dividing cells (left) or extruding cells (right). Relative cell areas are aligned according to the time of cytokinesis termination (left) or closure of apical area (right) corresponding to time 0. Orange lines are for uncorrected movies, and magenta lines after segmentation correction. Grey area is the confidence interval at 0.95.

**Figure Sup 5:**
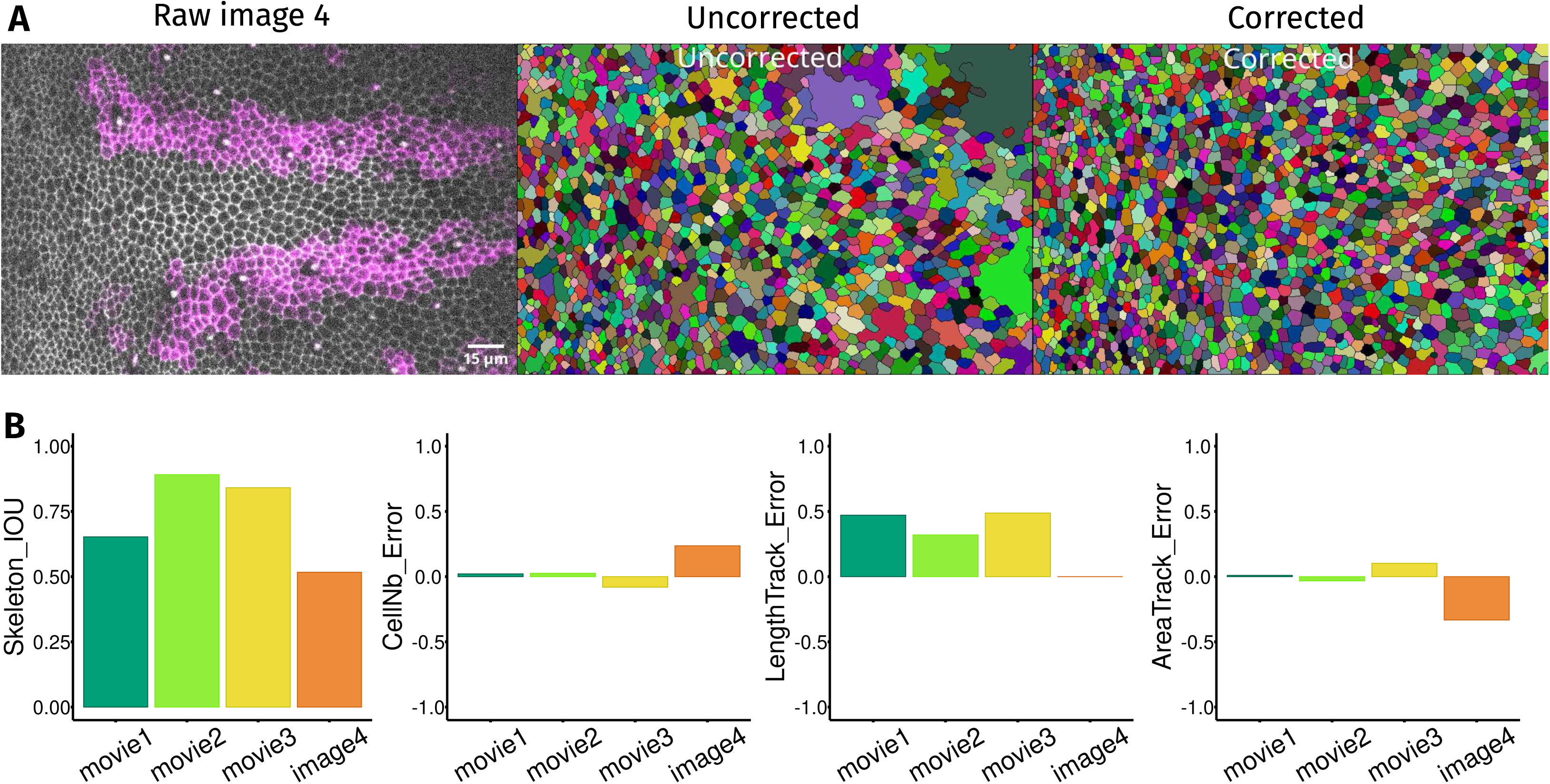
Segmentation error estimation for the four biological model systems. (A) Raw image of a *Drosophila* notum expressing E-cad::GFP (grey) with cells expressing RFP in the stripe-Gal4 domain (magenta) (left), segmentation before (middle) and after (right) correction of the 4^th^ example of Figure 5, called “Image4”. Scale bar is 15µm. (B) Estimation of the segmentation accuracy (the skeleton binary files before and after correction are compared, measuring the intersection-over-union (IOU) of the binaries), cell number error, length track error and area track error in the four model systems (see Figure 5). Error is defined as the relative difference between the measure after and before correction as: (after_correction – before_correction)/after_correction.

## Movie legends

**Movie S1:** EpiCure interface and display options. The full Napari interface is recorded while changing the display options of EpiCure.

**Movie S2:** Movie of the *Drosophila* pupal notum from 20 to 32 hours after pupal formation (left), with cell contours marked with E-Cad::GFP (grey). Cell segmentation (one color=one cell) before (middle) and after (right) correction. Scale bar=30μm.

**Movie S3:** Movie of the *Drosophila* pupal notum from 20 to 32 hours after pupal formation (left), with cell contours marked with E-Cad::GFP (grey). Detection of cellular events and marking of cell fate: cells that will divide later (blue) or extrude (red) for uncorrected segmentation (left) and after segmentation curation (right). The SOs (Sensory Organs) are shown in green. Scale bar=30μm.

**Movie S4:** Movie of quail gastrula extraembryonic tissue showing cytoplasmic membrane GFP (grey, left) with segmentation before (middle) and after (right) correction. Scale bar=15μm.

**Movie S5:** Movie of *Drosophila* abdominal histoblasts with contours marked with E-cad::GFP (grey, left) with segmentation before (middle) and after (right) correction. Scale bar=15μm.

**Movie S6:** Movie of the adult neural stem cell layer of the zebragish telencephalon with contour marked with gpap:Zo1::mKate2 (grey) and deltaA:eGFP (orange) (left) with segmentation before (middle) and after (right) correction. Scale bar=15μm.

